# Evolutionary arms race between virus and host drives genetic diversity in bat SARS related coronavirus spike genes

**DOI:** 10.1101/2020.05.13.093658

**Authors:** Hua Guo, Bing-Jie Hu, Xing-Lou Yang, Lei-Ping Zeng, Bei Li, Song-Ying Ouyang, Zheng-Li Shi

## Abstract

The Chinese horseshoe bat (*Rhinolophus sinicus*), reservoir host of severe acute respiratory syndrome coronavirus (SARS-CoV), carries many bat SARS-related CoVs (SARSr-CoVs) with high genetic diversity, particularly in the spike gene. Despite these variations, some bat SARSr-CoVs can utilize the orthologs of human SARS-CoV receptor, angiotensin-converting enzyme 2 (ACE2), for entry. It is speculated that the interaction between bat ACE2 and SARSr-CoV spike proteins drives diversity. Here, we have identified a series of *R. sinicus* ACE2 variants with some polymorphic sites involved in the interaction with the SARS-CoV spike protein. Pseudoviruses or SARSr-CoVs carrying different spike proteins showed different infection efficiency in cells transiently expressing bat ACE2 variants. Consistent results were observed by binding affinity assays between SARS- and SARSr-CoV spike proteins and receptor molecules from bats and humans. All tested bat SARSr-CoV spike proteins had a higher binding affinity to human ACE2 than to bat ACE2, although they showed a 10-fold lower binding affinity to human ACE2 compared with their SARS-CoV counterpart. Structure modeling revealed that the difference in binding affinity between spike and ACE2 might be caused by the alteration of some key residues in the interface of these two molecules. Molecular evolution analysis indicates that these residues were under strong positive selection. These results suggest that the SARSr-CoV spike protein and *R. sinicus* ACE2 may have coevolved over time and experienced selection pressure from each other, triggering the evolutionary arms race dynamics. It further proves that *R. sinicus* is the natural host of SARSr-CoVs.

**Importance:** Evolutionary arms race dynamics shape the diversity of viruses and their receptors. Identification of key residues which are involved in interspecies transmission is important to predict potential pathogen spillover from wildlife to humans. Previously, we have identified genetically diverse SARSr-CoV in Chinese horseshoe bats. Here, we show the highly polymorphic ACE2 in Chinese horseshoe bat populations. These ACE2 variants support SARS- and SARSr-CoV infection but with different binding affinity to different spike proteins. The higher binding affinity of SARSr-CoV spike to human ACE2 suggests that these viruses have the capacity of spillover to humans. The positive selection of residues at the interface between ACE2 and SARSr-CoV spike protein suggests a long-term and ongoing coevolutionary dynamics between them. Continued surveillance of this group of viruses in bats is necessary for the prevention of the next SARS-like disease.

## Introduction

Coronaviruses belong to the *Orthocoronavirinae* subfamily and *Coronaviridae* family. They are enveloped viruses and contain a positive-sense and single-stranded RNA genome. There are four genera in this subfamily, *Alphacoronavirus, Betacoronavirus, Gammacoronavirus*, and *Deltacoronavirus. Alphacoronavirus* and *Betacoronavirus* are believed to have originated in bats or rodents while *Gammacoronavirus* and *Deltacoronavirus* in birds (1, 2). Since the beginning of the 21^st^ century, three betacoronaviruses have caused outbreaks of severe pneumonia in humans. These are, the severe acute respiratory syndrome coronavirus (SARS-CoV) (3-5), the Middle-East respiratory syndrome coronavirus (MERS-CoV) (6) and the ongoing 2019 novel coronavirus (SARS-CoV-2) (7).

The SARS-CoV-2 outbreak has brought back memories of SARS-CoV that occurred 17 years ago in China. SARS is a zoonosis, as demonstrated by the identification of an almost identical coronavirus present in market civets in Guangdong province, China (8). In the following years, genetically diverse SARS-related coronaviruses (SARSr-CoV) were detected or isolated from horseshoe bats from different regions of China and Europe (9-18). Bat SARSr-CoVs share 78– 96% nucleotide sequence identity with human and civet SARS-CoVs, with the most variable regions encoding the spike protein (S) and accessory protein ORF3 and 8 (17, 19). Moreover, we have identified all the building blocks of SARS-CoV in the genome of different bat SARSr-CoVs and suggest that the ancestor of SARS-CoV originated in bats through the recombination of bat SARSr-CoV genomes (17, 19).

The first and essential step of virus infection is cell receptor recognition. The entry of the coronavirus is mediated by specific interactions between the viral S protein and cell surface receptor, followed by fusion between the viral and host membrane. The coronavirus S protein is functionally divided into two subunits, a cell attachment subunit (S1) and a membrane-fusion subunit (S2). The S1 region contains an N-terminal domain (NTD) and a C-terminal domain (CTD); both can be used for coronavirus receptor binding (RBD) (20). For SARS-CoV, its S1-CTD serves as an RBD for binding to the cellular receptor, angiotensin-converting enzyme 2 (ACE2) (21). Biochemical and crystal structure analyses have identified a few key residues in the interface between the SARS-CoV S-RBD and human ACE2 (21-23).

SARSr-CoVs, detected in *Rhinolophus sinicus*, can be divided into two distinct clades based on the size of the S protein. Clade 1 includes viruses that have S proteins identical in size to the SARS-CoV S protein, whereas viruses belonging to clade 2 have a smaller S protein, due to 5, 12, or 13 amino acid deletions (17, 19). Despite the variations in the RBD, all clade 1 strains can use ACE2 for cell entry, whereas clade 2 strains, with deletions cannot (14, 16, 17). These results suggest that members of clade 1 are likely to be the direct source of SARS-CoV in terms of genome similarity and ACE2 usage.

ACE2 is functionally divided into two domains—the N-terminal domain is involved in SARS-CoV binding and the C-terminal domain in the regulation of heart function (21, 24). Previous results have indicated that the C-terminal domains of ACE2 from different origins are relatively well conserved, whereas the N-terminal domains show much more diversity across species (25). Previously, we have shown that SARS-CoV can utilize ACE2 derived from *Myotis daubentonii* and *R. sinicus*; minor mutations in the RBD-binding site could convert ACE2 from being unsusceptible to susceptible to SARS-CoV binding (26, 27). As all bat SARSr-CoV belonging to clade 1 can utilize ACE2 and have been isolated from *R. sinicus*, we asked whether variations in ACE2 of *R. sinicus* could contribute to the diversity of bat SARSr-CoV.

Here, we have investigated the polymorphism of *R. sinicus* ACE2 genes and assessed their susceptibility and binding affinity to different bat SARSr-CoV spike proteins through a combination of molecular evolution analyses, protein affinity assays, and virus infection assays. Our results showed that the diversity of the SARSr-CoV spike protein may experience natural selection pressure from the *R. sinicus* ACE2 variants; over long periods of co-existence, the SARSr-CoV spike protein may be selected by *R. sinicus* ACE2 to maintain genetic diversity and to fit with the population of *R. sinicus*.

## Results

### ACE2 genes show high polymorphism among the *R. sinicus* populations

Samples from three provinces (Hubei, Guangdong, and Yunnan) were used for ACE2 amplification, based on the prevalence of bat SARSr-CoVs and tissue sample availability and quality. In addition to previously sequenced bat ACE2 by our group (sample ID 832, 411, and 3357, collected from Hubei, Guangxi, and Yunnan, respectively) and others (GenBank accession no. ACT66275; sample collected from Hong Kong), we obtained ACE2 gene sequences from 21 *R. sinicus* bat individuals: five from Hubei, nine from Guangdong, and seven from Yunnan. The ACE2 sequences exhibited 98–100% amino acid (aa) identity within their species and 80-81% aa identity with human ACE2 (**Table S1**). Major variations were observed at the N-terminal region, including in some residues which were previously identified to be in contact with SARS-CoV S-RBD (**Fig. 1A and Fig. S1**). Analysis based on nonsynonymous SNPs helped identify eight residues, including 24, 27, 31, 34, 35, 38, 41, and 42. The combination of these 8 residues produced eight alleles, including RIESEDYK, LIEFENYQ, RTESENYQ, RIKSEDYQ, QIKSEDYQ, RMTSEDYQ, EMKTKDHQ, and EIKTKDHQ, named allele 1–8, respectively (**Fig. 1A**). In addition to the ACE2 genotype data from previous studies (allele 4, 7, and 8), five novel alleles were identified in the *R. sinicus* populations in this study. Alleles 2 and 4 were found in two and three provinces, respectively, whereas the other alleles seemed to be geographically restricted. In summary, three alleles (4, 6, and 8) were found in Guangdong, four (1, 2, 4, and 7) in Yunnan, three (2, 4, and 5) in Hubei, and one each in Guangxi and Hong Kong. Coexistence of four alleles was found in the same bat cave of Yunnan where the direct progenitor of SARS-CoV was found (**Fig. 1B**). Taken together, these data suggest that ACE2 variants have been circulating within the *R. sinicus* populations in different regions for a long time; substitutions at the sites directly in contact with SARS-CoV S-RBD suggest that they may have important functions during SARS-CoV evolution and transmission.

**Fig. 1.**
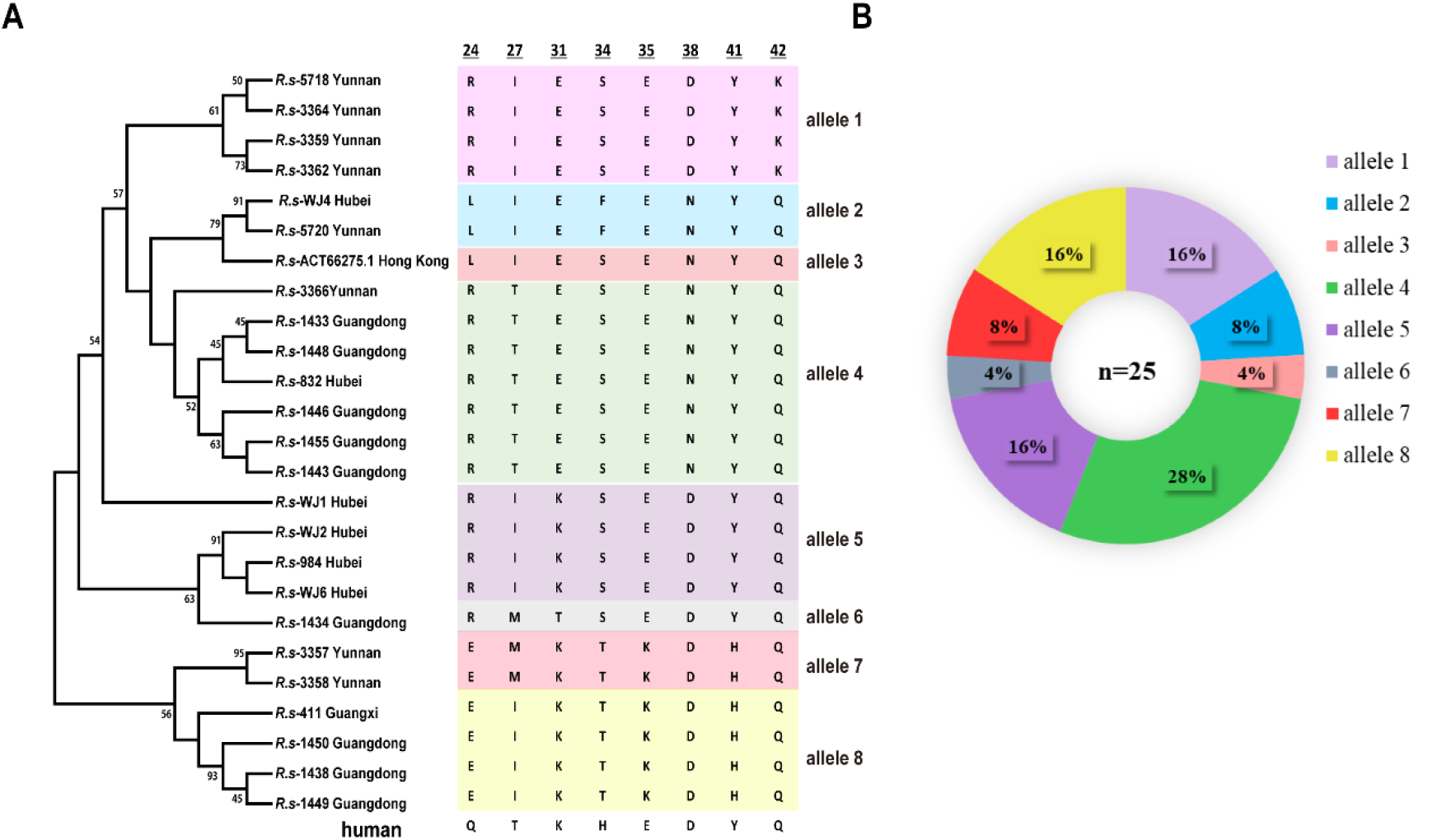
Phylogenetic tree of *R. sinicus* ACE2. (A) The maximum-likelihood tree (left panel) was produced using MEGA6 software, based on the alignment of ACE2 amino acid sequences of *R. sinicus* with the Jones-Taylor-Thornton (JTT+G+I) model and bootstrap value (%) of 1000 replicates (28). The eight key residues which are involved in interacting with the SARS-CoV spike are indicated in the panel on the right. The eight residues of human ACE2 are indicated at the bottom. The numbers at the top are amino acid positions in ACE2. (B) Frequencies of ACE2 alleles among the *R. sinicus* population. The number of *R. sinicus* ACE2 sequences is shown in the center of the pie chart. Different colored sectors with percentages in the pie chart indicate allele frequencies of ACE2 in the *R. sinicus* population used in this study. The alignments of complete amino acid sequences of *R. sinicus* ACE2 are shown in supplementary **Fig. S1**. ACE2 sequences of *Rs*-411, 832, and 3357 and *Rs*-ACT66275.1 have been published previously and were downloaded from GenBank. The accession number of *R. sinicus* ACE2 sequences obtained in this study are listed in Table S2

### *R. sinicus* ACE2 variants show different susceptibility to SARSr-CoV infection

To assess if different *R. sinicus* ACE2 molecules affect the entry of SARS-CoV and bat SARSr-CoV, *R. sinicus* ACE2 variants were transiently expressed in HeLa cells and the entry efficiency of SARS-CoV pseudotyped or bat SARSr-CoVs carrying different S proteins were tested. The four tested bat SARSr-CoV strains can be divided into four genotypes according to their S1 sequences, as reported previously (17). Briefly, compared with SARS-CoV S protein, SARSr-CoV-RsWIV1 shares high aa identity at the RBD; RsWIV16 is the closest relative of SARS-CoV and shows high aa identity at both the NTD and RBD; Rs4231 shares high aa identity at the NTD; and RsSHC014 shows divergence at both the NTD and RBD regions.

Similar to our previous report, all four bat SARSr-CoV strains with the same genomic background but different S proteins could use human ACE2 and replicate at similar levels (17). However, there are some differences in how they utilize *R. sinicus* ACE2s (**Fig. 2 and Fig. S2**). All test viruses could efficiently use allele 1, 2, 4, 5 for entry. RsWIV1 and RsWIV16, which share an identical RBD, could not use allele 6 (sample ID 1434) from Guangdong. Rs4231 and RsSHC014, which share an identical RBD, could not use allele 7 (sample ID 3357) and 8 (sample ID 1438) from Yunnan and Guangdong, respectively. SARS-CoV-BJ01, which shares high similarity with WIV1 and WIV16 RBD, was able to use same bat ACE2 alleles as Rs4231 and RsSHC014 in the pseudotyped infection assay (**Fig. 2 and Fig. S3**). These results indicate that cell entry was affected by both spike RBD and *R. sinicus* ACE2 variants.

**Fig. 2.**
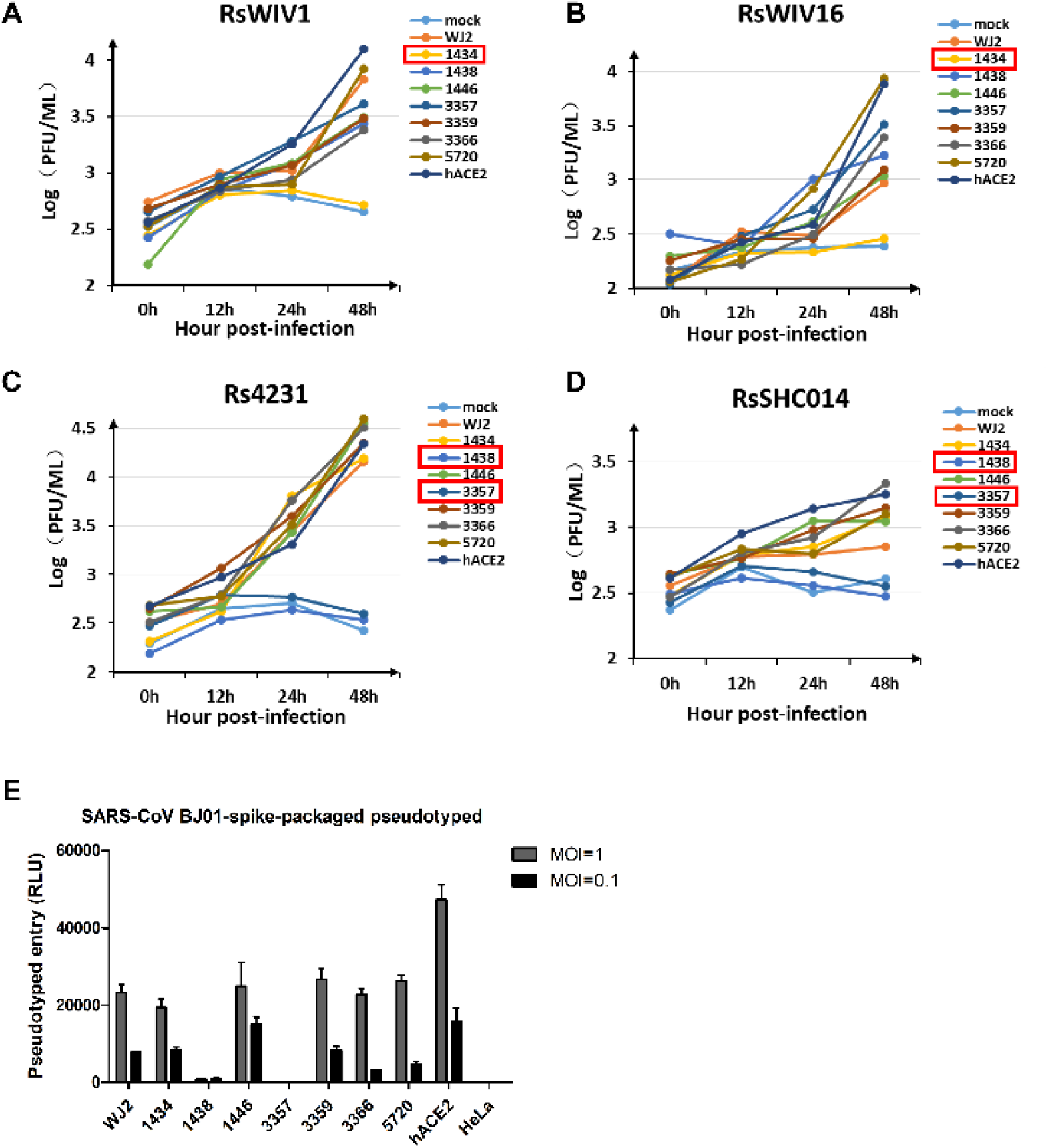
Infectivity of SARSr-CoV in HeLa cells expressing *R. sinicus* ACE2. (A–D) Determination of bat SARSr-CoV infectivity in HeLa cells with and without the expression of ACE2 from *R. sinicus* or human (hACE2) at an MOI = 0.01. The growth curves were determined by real-time PCR. The number in a red square indicates the sample which is not susceptible to bat SARSr-CoV. (E) The infectivity of the SARS-CoV-BJ01 pseudotyped was used for the assay at an MOI = 1 and 0.1, due to biosafety regulation in China. Error bars represent the SEM from two independent transfections; each assay was performed in triplicate.

### Mutation of *R. sinicus* ACE2 residues affects its binding affinity with SARSr-CoV RBDs

To further explain the different ability of SARS and SARSr-CoVs in ACE2 usage, we expressed RBD from SARS-CoV BJ01, RsWIV1, and RsSHC014 and three *R. sinicus* ACE2s, allele 6 (sample 1434), allele 7 (sample 3357), and allele 2 (sample 5720), and tested the binding affinity between them. BJ01 RBD with human ACE2 and RsWIV1 RBD with human DPP4 were used as the positive and negative control, respectively. Real-time analysis showed that the binding affinity between different RBDs and ACE2s was different based on the equilibrium dissociation constant (KD) (**Fig. 3 and Fig. S4**). Consistent with the results of the virus infection experiments, 1434ACE2 (allele 6) was found to bind RsSHC014 and BJ01 but not RsWIV1 RBD; 3357ACE2 (allele 7) was found to bind RsWIV1 but not RsSHC014 and BJ01 RBD; 5720ACE2 (allele 2) was found to bind all tested RBDs. All tested RBDs had a high binding affinity to human or bat ACE2. BJ01 RBD had a higher binding affinity for human ACE2 than did RsWIV1 and RsSHC014 RBDs (**Fig. 3A, E, and I**); however, it had a lower binding affinity to bat ACE2 than the two bat SARSr-CoV RBDs (**Fig. 3B, D, F, G, J, K**). These results demonstrated that the binding affinity between spike RBD and ACE2 affects the ability of the virus to infect.

**Fig. 3.**
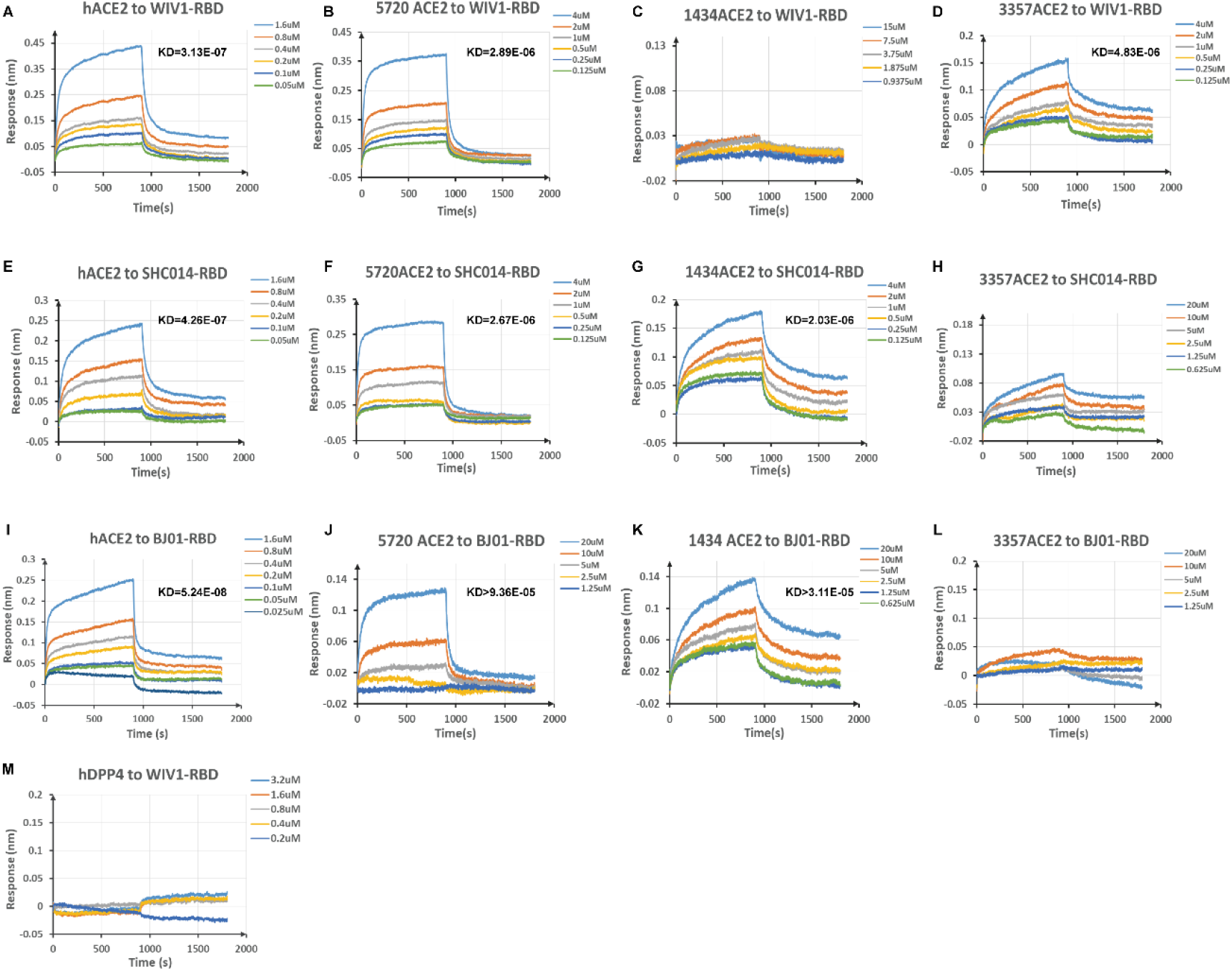
Binding affinity assay between different RBDs and ACE2s. (A–D) Binding assay of human ACE2 or bat ACE2 to WIV1 RBD. (E–H) Binding assay of human ACE2 or bat ACE2 to SHC014 RBD. (I–L) Binding assay of human ACE2 or bat ACE2 to BJ01 RBD. (M) Binding assay of human DPP4 to WIV1 RBD was used as the negative control. The parameters of KD value (M) are shown on the upper right side of the picture. Different RBD proteins were immobilized on the sensors and tested for binding with graded concentrations of *R. sinicus* ACE2s, hACE2, or hDPP4. The Y-axis shows the real-time binding response.

### Structure modeling of the interaction between SARSr-CoV RBDs and *R. sinicus* ACE2s

The four tested spike proteins of bat SARSr-CoV are identical in size and share over 90% aa identity with SARS-CoV, which suggests that these proteins have a similar structure. In this study, we built structural complex models of bat SARSr-CoV-RsWIV1 RBD with *R. sinicus* ACE2 3357 (allele 7) and RsSHC014 RBD with *R. sinicus* ACE2 1434 (allele 6), in concordance with the results of the binding affinity assay between SARS-CoV RBD and human ACE2 (**Fig. 4**). Compared with the contact residues in the interface between SARS-CoV RBD and human ACE2, changes in or near the two virus-binding hotspots on ACE2 (hotspots Lys31 and Lys353 consist of a salt bridge with Glu35 and Asp38, respectively) were observed. As reported previously, the two hot spots are buried in a hydrophobic environment. They provide a substantial amount of energy to the virus-receptor binding interactions as well as fill critical voids in the hydrophobic stacking interactions at the binding interface (21, 29).

**Fig. 4.**
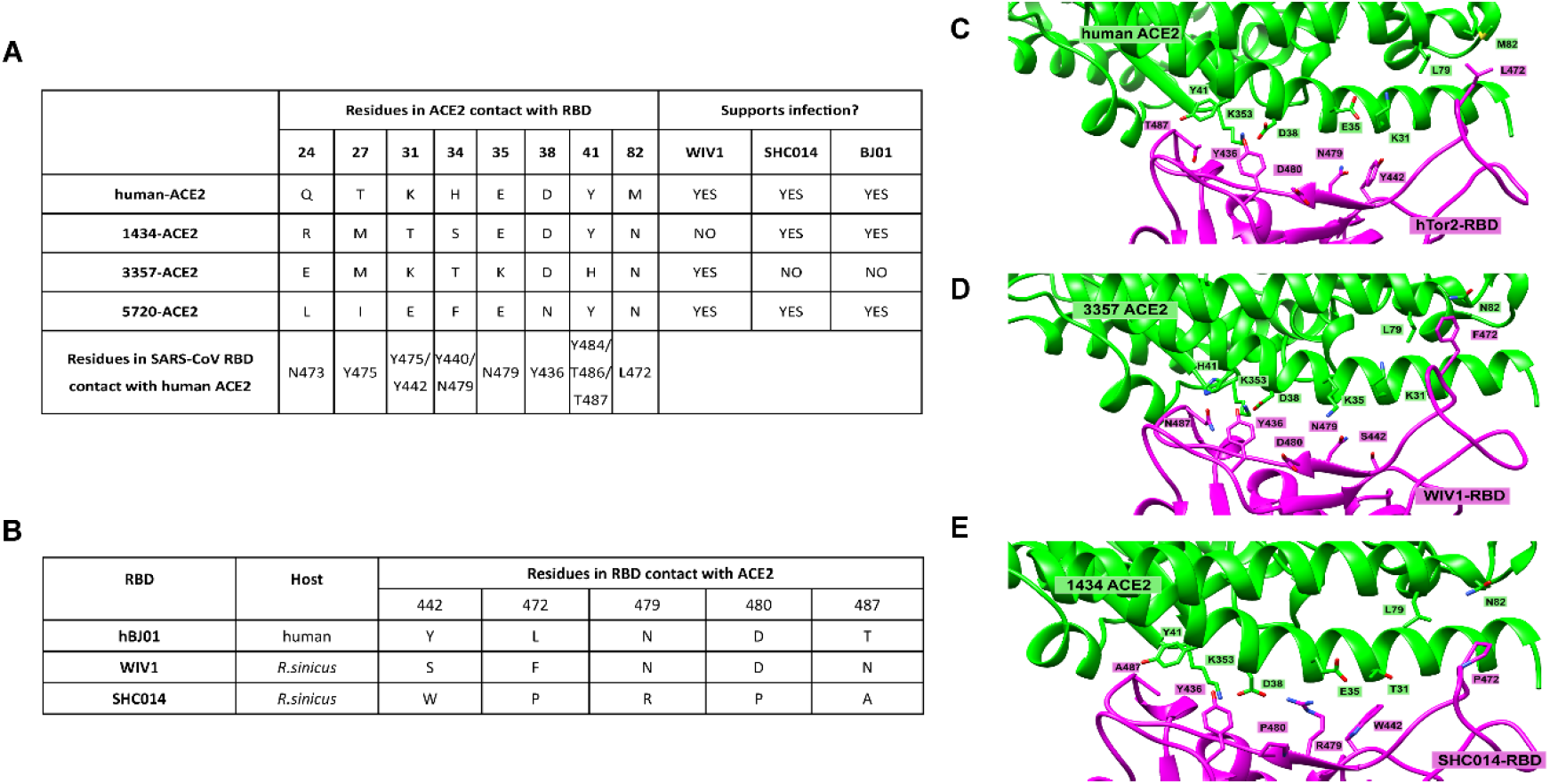
Structure modeling at the interface between SARSr-CoV spike and human or bat ACE2. Detailed view of the interaction between RBD and ACE2. Several important residues in RBD and ACE2 that are involved in the interactions are shown. ACE2 residues are in green and RBD residues are in magenta. (A) List of eight contact residues in ACE2 from human and *R. sinicus* that are directly involved in RBM/ACE2 binding. (B) Alignment of five residues from SARS-CoV and two SARSr-CoVs RBD that are critical for receptor binding. (C) Structure complex of the SARS-CoV hTor2 RBD and human ACE2 (protein data code 2AJF). (D) Predicted structure complex of *R. sinicus* ACE2 3357 with RsWIV1 RBD. (E) Predicted structure complex of *R. sinicus* ACE2 1434 with RsSHC014 RBD. The model for RsWIV1 RBD and ACE2 was built based on the structure of hTor2 RBD with hACE2 (protein data code 2AJF). The model for RsSHC014 RBD and ACE2 was built based on the structure of civet-optimized RBD with hACE2 (protein data code 3CSJ). The model for (C) has been adapted from previous studies (19, 29).

Compared with human ACE2 (**Fig. 4A, C, D**), *R. sinicus* ACE2-3357 contains two main residue changes, E35K and Y41H. The E35K breaks the salt bridge with K31 and Arg479 in RsSHC014 RBD, which may influence the binding affinity between them. A histidine at position 41 may weaken the support for the K353–D38 salt bridge because it is a weaker hydrophobic stacker than tyrosine, resulting in decreased binding affinity with BJ01 RBD (30). Consequently, BJ01 and RsSHC014 RBD showed a lower binding affinity with *R. sinicus* ACE2-3357.

In *R. sinicus* ACE2-1434, we found a threonine at 31, unlike human ACE2, which has a lysine at this position (**Fig. 4**). Although the K31T change would fail to form a salt bridge with Glu35, Tyr442 in BJ01 RBD can form a hydrogen bond with it (29). However, a serine at 442 in RsWIV1 RBD cannot. Besides, RsSHC014 contains an arginine at 479, and Thr31 cannot form a salt bridge with Glu35, making Glu35 available to form a salt bridge with Arg479, but the RBD residue Asn479 in RsWIV1 may lose this ability. Consequently, BJ01 and RsSHC014 RBD can bind with *R. sinicus* ACE2-1434, but not RsWIV1.

In addition, all *R. sinicus* ACE2s in this study contain an asparagine at position 82 rather than a methionine in human ACE2. The M82N change introduces an unfavorable hydrophilic residue which would weaken the hydrophobic network around hot spot 31. Moreover, Asn82 introduces a glycosylation site, like that in rat ACE2, which cannot support SARS-CoV infection efficiently; the glycan at position 82 of ACE2 may lead to steric interference with viral RBD binding. Hence, M82N may have significant effects on the interaction between SARS-CoV and SARSr-CoV RBD with *R. sinicus* ACE2s. As previously described, residue 487 of the RBD interacts with hot spot 353 on ACE2. RsWIV1 contains an Asn487 in its RBD, the polar side chain of Asn487 may have unfavorable interactions with the aliphatic portion of residue Lys353 in ACE2, which may affect the hot spot interaction between K353 and D38. Moreover, the RsSHC014 RBD contains an alanine at position 487; the small side chain of Ala 487 does not provide support to the structure of hot spot 353 (19). Therefore, both RsWIV1 and RsSHC014 RBD had a lower binding affinity to human ACE2 than did BJ01, but they both showed a higher binding affinity with human ACE2 than *R. sinicus* ACE2s, which correlates well with the results from virus infection and binding assays.

### SARSr-CoV spike genes have coevolved with *R. sinicus* ACE2 through positive selection

To test the possible selection pressure acting on the SARSr-CoV spike and *R. sinicus* ACE2 gene, we used the codeml program of the PAML software package (31) to analyze the ratio of nonsynonymous to synonymous mutations (dN/dS ratios) at individual codons. We analyzed the complete gene encoding the SARSr-CoV spike protein by aligning nine SARSr-CoV spike gene sequences from *R. sinicus* samples. We found that the models allowed codons to evolve under positive selection (M2a and M8). In model M8 (initial seed value for ω (dN/dS) = 1.6, codon frequency = F3×4), 20 codons (p>0.95) were under positive selection with dN/dS>1 (**Table 1**), 17 of those were found to be located on the RBD region, which faces its receptor ACE2, according to the crystal structure (**Fig. S5)**. Moreover, five of those (442, 472, 479, 480, and 487), present in the SARS-CoV spike, have been previously identified to have a significant impact on binding affinity to human ACE2 (**Fig. 4, Table S3 and Fig. S5**) (32). We next analyzed the ACE2 gene of *R. sinicus* by aligning 25 *R. sinicus* ACE2 gene sequences obtained in this study and downloaded from GenBank. We found that 12 codons (2.3%, p>0.95) were under positive selection with dN/dS>1 in model M8 (**Table 1**), and 8 of them (24, 27, 31, 34, 35, 38, 41, and 42) correspond to the residues in human ACE2, which were previously identified to be involved in direct contact with the human SARS-CoV spike protein (**Fig. 4, Fig. S5**) (32). We also analyzed the ACE2 gene of *Rhinolophus affinis* (*R. affinis*), which has been reported to carry SARSr-CoV occasionally (15). Used an alignment of 23 ACE2 gene sequences from *R. affinis* obtained in this study, we found that *R. affinis* ACE2 was more conserved between different individuals in the entire coding region than *R. sinicus* ACE2 (**Fig. S6**) and no obvious positive selection sites were observed (data not shown). Additionally, by querying single nucleotide polymorphism (SNP) databases, we found that SNPs in human ACE2 gene arose randomly through the entire coding region (https://www.ncbi.nlm.nih.gov/snp/). Although we identified two SNP sites with nonsynonymous mutation (T27A and E35K) in human ACE2, both displayed a rare frequency in the global population (frequency 0.00001 and 0.00002, respectively). These results indicate that positive selection has happened at the interface between bat SARSr-CoV spike protein and *R. sinicus* ACE2.

**Table 1.** PAML analysis of *R. sinicus* ACE2 and SARSr-CoV spike sequences.

| dataset | $\omega$ 0, | Model comparison <sup>c</sup> | | | | | | dN/dS | Residues with dN/dS>1 |
| --- | --- | --- | --- | --- | --- | --- | --- | --- | --- |
| <sup>a</sup> | codon | M1a vs M2a |  | M7 vs M8 |  | M8a vs M8 |  | (%)<sup>d</sup> |  |
|  | freq<sup>b</sup> |  |  |  |  |  |  |  | *p>0.95 **p>0.99<sup>e</sup> |
| | | 2δ | p-value | 2δ | p-value | 2δ | p-value | | |
| ACE2 | 0.4, F61 | 70.92 | p=4.441E-16 | 60.66 | p=6.728E-14 | 11.82 | p=1.356E-3 | 16.5(1.9) | S5*,L24**,I27**,E31**,F34**,E35*,N38** |
| Full |  |  |  |  |  |  |  |  | ,Y41*,Q42*,P84**,H139*,N159** |
| length | 0.4, F3X4 | 72.26 | p=2.22E-16 | 74.32 | p=1.11E-16 | 72.27 | p=1.11E-16 | 15.6(2.3) | S5*,L24**,I27**,E31**,F34**,E35*,N38** |
|  |  |  |  |  |  |  |  |  | ,Y41*,Q42*,P84**,H139*,N159** |
|  | 1.6, F61 | 70.92 | p=4.441E-16 | 74.58 | p=1.11E-16 | 11.82 | p=1.356E-3 | 16.5(1.9) | S5*,L24**,I27**,E31**,F34**,E35*,N38** |
|  |  |  |  |  |  |  |  |  | ,Y41*,Q42*,P84**,H139*,N159** |
|  | 1.6, F3X4 | 72.26 | P=2.22E-16 | 74.32 | p=1.11E-16 | 72.27 | p=1.11E-16 | 15.4(2.3) | S5*,L24**,I27**,E31**,F34**,E35*,N38** |
|  |  |  |  |  |  |  |  |  | ,Y41*,Q42*,P84**,H139*,N159** |
| Spike | 0.4, F61 | 59.52 | p=1.189E-13 | 80.54 | p<1.11E-16 | 59.52 | p=5.945E-14 | 3.9(4.9) | Y151*,T417**,R426**,N427**,I428*,Q43 |
| Full |  |  |  |  |  |  |  |  | 2**,K439**,S442*,H445**,G446**,R449* |
| length |  |  |  |  |  |  |  |  | *,I455*,V458*,P459**,D463*,P470**,F47 |
|  |  |  |  |  |  |  |  |  | 2**,W476**,N479**,D480**,N487** |
|  | 0.4, F3X4 | 54.22 | p=1.684E-12 | 61.86 | p=4.041E-14 | 49.86 | p=7.445E-12 | 7.6(2) | H37*,Y151*,S198*,T417**,R426**,N427* |
|  |  |  |  |  |  |  |  |  | **,Q432**,K439**,S442*,H445**,G446** |
|  |  |  |  |  |  |  |  |  | ,R449**,V458*,P459*,P470*,F472*,W476 |
|  |  |  |  |  |  |  |  |  | **,N479*,D480**,N487* |
|  | 1.6, F61 | 59.52 | p=1.189E-13 | 80.54 | p<1.11E-16 | 59.52 | p=5.945E-14 | 3.9(4.9) | Y151*,T417**,R426**,N427**,I428*,Q43 |
|  |  |  |  |  |  |  |  |  | 2**,K439**,S442*,H445**,G446**,R449* |
|  |  |  |  |  |  |  |  |  | *,I455*,V458*,P459**,D463*,P470**,F47 |
|  |  |  |  |  |  |  |  |  | 2**,W476**,N479**,D480**,N487** |
|  | 1.6, F3X4 | 54.22 | p=1.684E-12 | 61.86 | p=4.041E-14 | 49.86 | p=7.445E-12 | 7.6(2) | H37*,Y151*,S198*,T417**,R426**,N427* |
|  |  |  |  |  |  |  |  |  | **,Q432**,K439**,S442*,H445**,G446** |
|  |  |  |  |  |  |  |  |  | ,R449**,V458*,P459*,P470*,F472*,W476 |
|  |  |  |  |  |  |  |  |  | **,N479*,D480**,N487* |

## Discussion

Chinese horseshoe bats are widely distributed in China and carry genetically diverse SARSr-CoVs. Two different clades of bat SARSr-CoVs were discovered, based on the size and similarity of S protein to that of human SARS-CoV. SARSr-CoVs of clade-1, only found in Yunnan province, have an S protein identical in size to SARS-CoV and use the ACE2 receptor, whereas SARSr-CoVs of clade-2 are widely distributed in China and cannot use ACE2 as the receptor (19). In this study, we analyzed ACE2 sequences of *R. sinicus*, collected from four provinces and one city, and observed highly polymorphic sites, which correspond to those that interact with SARS-CoV RBD, at the N-terminal region. Despite these variations, most ACE2s supported the viral entry of clade-1 bat SARSr-CoVs, but with different susceptibilities and binding affinities. In addition, we have identified key residues involved in the interaction between the bat ACE2 variants and SARSr-CoV RBD by structural modeling and positive selection analysis. These results indicate that SARSr-CoV has co-evolved with its natural host, *R. sinicus*, for a very long time. Members of clade-1 use ACE2 as the receptor, whereas members of clade-2 exploit different receptor(s), due to the deletions in S protein.

Genes with important functions usually display a dN/dS ratio of less than 1 (negative selection) because most amino acid alterations in a protein are deleterious. In a host-virus arms race situation, the genes involved tend to display dN/dS ratios greater than 1 (positive selection), specifically in the codons involved in the interaction interface between the virus and its host, with minimal effect on their physical function (20, 33). In this study, our analysis of the SARSr-CoV spike protein and *R. sinicus* ACE2 gene sequences showed that some codons related to the interaction interface between them were under positive selection. Similar rapid adaptation for SARS-CoV occurred during the SARS outbreak in 2002–2003 (21, 34). When SARS-CoV was transmitted from market civet to human, the spike gene experienced positive selection, where mutations in two critical residues (amino acids 479 and 487) of the spike protein changed the binding affinity of the virus to human ACE2 from low to high; subsequently, turning into a pandemic strain (34).

An increasing number of studies have suggested that co-evolution is driven by specific interactions between high levels of virus sequence divergence and polymorphic host receptors (20, 33, 35-38). The first example was observed for the avian leucosis virus infection in which receptor polymorphism in chicken could alter the sensitivity to virus entry (36). Arms race can also occur between the glycoproteins of two different viruses, as seen for the rodent arenavirus and the mouse mammary tumor virus, with their receptor transferrin receptor 1 (TfR1) (39).

In a continuous coevolutionary process, viruses often tend to decrease their virulence to fit both the host and the virus population. The “trade-off hypothesis” is the most popular explanation for why pathogens often do not reach their maximum reproductive potential (40, 41). In this study, we have found that bat SARSr-CoV RBDs showed a lower binding affinity to *R. sinicus* ACE2 than to human ACE2, but did not display obvious differences among the *R. sinicus* ACE2s, indicating that bat SARSr-CoVs may decrease their virulence to fit both the host and themselves. When they adapt to fit other host receptors better, cross-species transmission events could happen. This situation can be demonstrated by the higher binding affinity seen between SARS-CoV-BJ01 RBDs and human ACE2s than bat SARSr-CoV strains in this study. Similar examples can be found in other cases of coevolution between host receptor and virus (36, 39, 42, 43). Previous studies have shown adaptive evolution of the coronavirus spike protein to host receptor *in vivo* and *in vitro* (44-48). Recently, the MERS-CoV spike protein was shown to rapidly adapt to *Desmodus rotundus* DPP4 (*dr*DPP4) receptor from semi-permissive to permissive state after several passages *in vitro*. Furthermore, mutations detected in the RBD of MERS-CoV spike protein enhanced viral entry and replication on cells expressing *dr*DPP4 within three passages (49). Similarly, adaptive mutations have occurred in the Ebola virus envelope protein, which contacts its receptor, NPC intracellular cholesterol transporter 1, when it crossed the host-virus barriers, was transmitted to different hosts, and entered the human population (50-53). These examples are a warning that the SARSr-CoVs circulating in *R. sinicus* may adapt to other animal hosts, including humans, and cause the next SARS-like disease. Therefore, it is important to continually monitor SARSr-CoVs in *R. sinicus* populations to avoid future spillover events. Our study provides an example to assess the risk of a potential spillover from viruses circulating in animal populations through the positive selection analysis, structural modeling, and experimental verification.

ACE2, a multifunctional enzyme involved in the negative regulation of the renin-angiotensin system, also interacts with amino acid transporters and integrins (54, 55). ACE2 functions as a carboxypeptidase and its role as the SARS-CoV receptor does not affect its peptidase activity. Moreover, the structural complex shows that the SARS-CoV spike RBD does not block the catalytically-active site of ACE2 (21, 56). Considering the conserved recognition site in receptor of SARSr-CoV clade 1, the interface of SARS-CoV and ACE2 interaction would be an ideal target for drug screening against a broad range of infections caused by these viruses.

## Materials and Methods

All work with the infectious virus was performed under biosafety level 2 conditions with appropriate personal protection.

### Bat samples, cells, and viruses

Bat samples were from the biobank of our laboratory. Vero E6, HeLa, and HEK 293T/17 cells (ATCC) were maintained in Dulbecco’s modified Eagle medium (DMEM) supplemented with 10% fetal bovine serum (FBS). Cells were cultured at 37°C with 5% CO_2_. Bat SARSr-CoV-RsWIV1 and recombinant viruses with RsWIV1 as the backbone and replaced by different spike genes of bat SARSr-CoVs (RsWIV16, RsSHC014, and Rs4231) were cultured as previously described (16, 57). Titers of all viruses used in this study were determined by plaque assays on Vero E6 cells, as previously described (17).

### Amplification, cloning, and expression of bats ACE2 gene

Bat ACE2 gene was amplified using DNA from the intestinal tissue of bats. In brief, total RNA was extracted from bat intestine tissue using the RNAprep pure Kit (for Cell/Bacteria) (TIANGEN, Beijing, China). First-strand complementary DNA was synthesized from total RNA by reverse transcription with random hexamers; full-length bat ACE2 fragments were amplified by reverse-transcription nested polymerase chain reaction (RT-PCR). Primers were designed based on available ACE2 sequences from NCBI. First-round primers are as follows: F-ACE2-out-AATGGGGTTTTGGCGCTCAG, R-ACE2-out-CATACAATGAAATCACCTCAAGAG, second-round primers: F-ACE2-in-CAGGGAAAGATGTCAGGCTC, R-ACE2-in-TTCTAAAABGAVGTYTGAAC. The ACE2 gene was cloned into the pcDNA3.1 vector with XhoI and BamHI. N-terminal mouse Igk or human IFNα1 signal peptide and S-tag were inserted. The plasmids were verified by sequencing. The expression of human and *R. sinicus* ACE2 was confirmed by western blotting and immunofluorescence assay.

### Expression constructs, protein expression, and purification

The constructs used for protein expression and purification were individually prepared by inserting coding sequences for SARS-CoV-BJ01 RBD (spike residues: aa 317–569, accession number: AY278488.2), RsWIV1 RBD (spike residues: aa 318–570, accession number: KF367457.1), RsSHC014 RBD (spike residues: aa 318–570, accession number: KC881005.1), *R. sinicus* ACE2 (aa 19–615, accession numbers are listed in Table S2), human ACE2 (aa 19–615, accession number: BAJ21180), and human DPP4 (aa 39–766, accession number: NP_001926) into the expression vector, as described previously (58). For each protein, an N-terminal signal peptide and an S-tag were added to facilitate protein secretion and purification. The proteins used for the Octet RED binding assay were expressed in HEK 293T/17 cells. Cells were washed twice with D-Hanks solution 6 h post-transfection, then cultured in fresh 293T FreeStyle expression medium. The cells were collected 48 h post-transfection and centrifuged at 4000 × *g* for 10 min at 4°C. The supernatant was incubated with S-tag agarose beads overnight at 4°C, passed through a column, and the protein was eluted from the column with 3 M MgCl_2_. The protein was finally buffered with PBS and stored at −80 °C until further use.

### Immunofluorescence assay

HeLa cells were seeded in 24-well plates and transfected with equal amounts of human or *R. sinicus* ACE2 plasmids. After 24 h, the cells were incubated with different SARSr-CoV strains at a multiplicity of infection (MOI) = 1 for 1 h. Thereafter, the cells were washed with D-Hanks solution thrice and supplemented with the medium. At 24 h after infection, cells were washed with PBS and fixed with 4% formaldehyde for 30 min at room temperature (about 25°C). ACE2 expression was detected using mouse anti-S-tag monoclonal antibody (Kyab Biotech Co. Ltd., Wuhan, China), followed by FITC-labelled goat anti-mouse IgG H&L (Abcam, Cambridge Biomedical Campus, Cambridge, UK). Virus replication was detected using rabbit serum against the SARSr-CoV-Rp3 NP, as previously described (14), and Cy3-conjugated goat anti-rabbit IgG (Abcam, Cambridge Biomedical Campus, Cambridge, UK). Nuclei were stained with 4′,6-diamidino-2-phenylindole. Staining patterns were examined using a fluorescence imaging system (EVOS(tm) FL Color Imaging System, Thermo Fisher Scientific, Waltham, MA, USA)

### Real-time PCR

SARSr-CoV infection was detected by RT-PCR, as described previously (14). In brief, HeLa cells were transfected with equal amounts of human or *R. sinicus* ACE2 plasmids, 24 h before they were infected with the virus, at an MOI = 0.01 with 1 h adsorption. Thereafter, the cells were washed with D-Hanks solution three times and cultured in 1 mL DMEM+2% FBS. The viral supernatants were harvested at 0, 12, 24, and 48 h post-infection—200 μL supernatant was removed and an equal amount of medium was added back at each time point. Viral RNA was extracted from the supernatant using the Viral RNA Mini Kit (Qiagen, Waltham, MA, USA) and then quantified on the ABI Step One system (Thermo Fisher Scientific, Waltham, MA, USA), with the AgPath-ID One-Step RT-PCR Kit (Applied Biosystems life technologies, Waltham, MA, USA). RNA dilutions from purified SARSr-CoV-RsWIV1 stock were used as a standard (with a titer of 6.5 × 10^6^ PFU/mL). Every sample was analyzed in triplicate on two independent occasions.

### Binding assay

Binding assays between SARS/SARSr-CoV RBD and human or *R. sinicus* ACE2 protein were performed using the Octet RED system (ForteBio, Menlo Park, CA, USA) in 96-well microplates at 30°C, as described previously (59, 60). Briefly, the RBD was biotinylated using EZ-Link NHS-LC-LC-Biotin (Thermo Fisher Scientific, Waltham, MA, USA). The assay was carried out by placing the Streptavidin Biosensors (ForteBio, Menlo Park, CA, USA) in the wells and measuring changes in layer thickness (nm) over time (s). First, the sensors were rinsed in the kinetic buffer (1 M NaCl, 0.1% BSA, 0.02% Tween-20; pH 6.5) for 120 s, which served as the baseline. Thereafter, the sensors were immobilized for 600 s, with 200 μL buffer containing biotinylated SARSr-CoV RBD (50 μg/mL). Subsequently, the sensors were washed in the kinetic buffer for 200 s. Finally, the sensors were exposed to a series of diluted ACE2 protein run in 200 μL volumes. The association between each RBD and its corresponding ACE2 binding partner was monitored for 900 s. This was followed by monitoring their dissociation in the kinetic buffer for 900 s. During the entire experiment, the sample plate was kept shaking at 1000 rpm, and an RBD-loaded biosensor was used in association with the buffer as a baseline. Data analysis from the ForteBio Octet RED instrument includes reference subtraction. Inter-step correction and Y-alignment were used to minimize tip-dependent variability. Data were globally fitted in a 1:1 model using the Data Analysis Software v7.1 (ForteBio, Menlo Park, CA, USA).

### Pseudotyped production and infection of ACE2-transfected HeLa cells

SARS-CoV-BJ01 pseudotyped particles were produced by HEK 293T/17 cells. Plasmids, pcDNA3.1-BJ01-S and pNL4-3.luc.R-E-, were co-transfected into HEK 293T/17 cells. At 6 h post-transfection, the medium was replaced with fresh 293T FreeStyle expression medium. The supernatant, which contained the pseudotyped particles, was harvested 48 h post-transfection and filtered using 0.45 μm filters.

Thereafter, the pseudotyped particles were aliquoted and stored at −80°C until further use. The viral titer was determined by the HIV p24 Quantitation ELISA Kits (Kyab Biotech Co. Ltd., Wuhan, China) before the viruses were used for the infection assay. HeLa cells were transfected with the same amount of human or *R. sinicus* ACE2 plasmids. After 24 h, the cells were incubated with SARS-CoV-BJ01 pseudotyped at an MOI = 1 or 0.1 for 1 h at 37°C, washed twice with D-Hanks solution, and supplemented with DMEM containing 10% FBS. Luciferase activity was determined using a GloMax luminometer (Promega Biotech Co. Ltd., Beijing, China) 48 h after infection. Each sample was analyzed in triplicate on two independent occasions.

### Structure modeling

The structure complex of SARS-CoV/SARSr-CoVs RBD and *R. sinicus* ACE2 was homology modeled using SWISS-MODEL (https://swissmodel.expasy.org/), based on the structure of human SARS-CoV RBD (hTor2 RBD) complexed with human ACE2 (hACE2) and civet-optimized RBD complexed with hACE2 (Protein Data Bank ID: 2AJF, 3SCJ) (29). Molecular graphics visualization and analyses were performed using the UCSF Chimera software (http://www.rbvi.ucsf.edu/chimera).

### Codon-based analysis of molecular evolution

Bat ACE2 and SARSr-CoV spike sequences were analyzed for positive selection. In this study, bat ACE2 sequences were either amplified or downloaded from NCBI and SARSr-CoV spike sequences were downloaded from NCBI; the database accession numbers are listed in Table S2. Sequences were aligned in Clustal X. Phylogenetic trees were built by the maximum likelihood method implemented in RAxML program in CIPRES Science Gateway (https://www.phylo.org/). Maximum likelihood analysis of dN/dS was performed using the codeml program in the PAML4.7 software package, as previously described (31, 39). In brief, multiple alignments were fit to NSsites models M1a, M2a, M7, M8a, and M8. Model fitting was performed with multiple seed values for dN/dS and assuming either the F61 or F3×4 model of codon frequencies. We used likelihood ratio tests (LRTs) to assess a better fit of codons that allowed positive selection. Posterior probabilities of codons under positive selection in different models were inferred using the Bayes empirical Bayes (BEB) algorithm.

### Accession numbers

The complete ACE2 sequences of *R. sinicus* and *R. affinis* ACE2 obtained in this study have been deposited in the GenBank database and the accession numbers are listed in Table S2.

## Acknowledgments

This work was jointly funded by the strategic priority research program of the Chinese Academy of Sciences (XDB29010101), National Natural Science Foundation of China (31770175 and 31621061) to Z.L.S.

We would like to thank Dr. Bing Yan for providing S-tag Agarose beads and mouse anti-S-tag monoclonal antibodies. We also thank Dr. Ding Gao in the Core Facility and Technical Support of the Wuhan Institute of Virology for his help in Octet RED technology. We thank Microorganisms & Viruses Culture Collection Center, Wuhan Institute of Virology, Chinese Academy of Sciences for providing HeLa and HEK 293T/17 cells.

## Declaration of Interests

The authors declare no competing interests.

## Author contributions

H.G. and Z.L.S. designed the research; H.G., B.J.H., L.P.Z., and B.L. performed the research; H.G. and Z.L.S. analyzed the data; H.G., X.L.Y., S.Y.O.Y., and Z.L.S. wrote the paper.

H.G. and B.J.H. contributed equally to this work.

## Notes

### Competing Interest Statement

The authors have declared no competing interest.

