## Supplementary material for "Evolutionary arms race between virus and host drives genetic diversity in bat SARS related coronavirus spike genes": Fig S1-S6, Table S2-3

**Fig. S1. Alignment of amino acids sequences of *R.sinicus* ACE2.** Residues 1-805 are shown. The accession numbers of various sequences used in this alignment are list in Table S2.

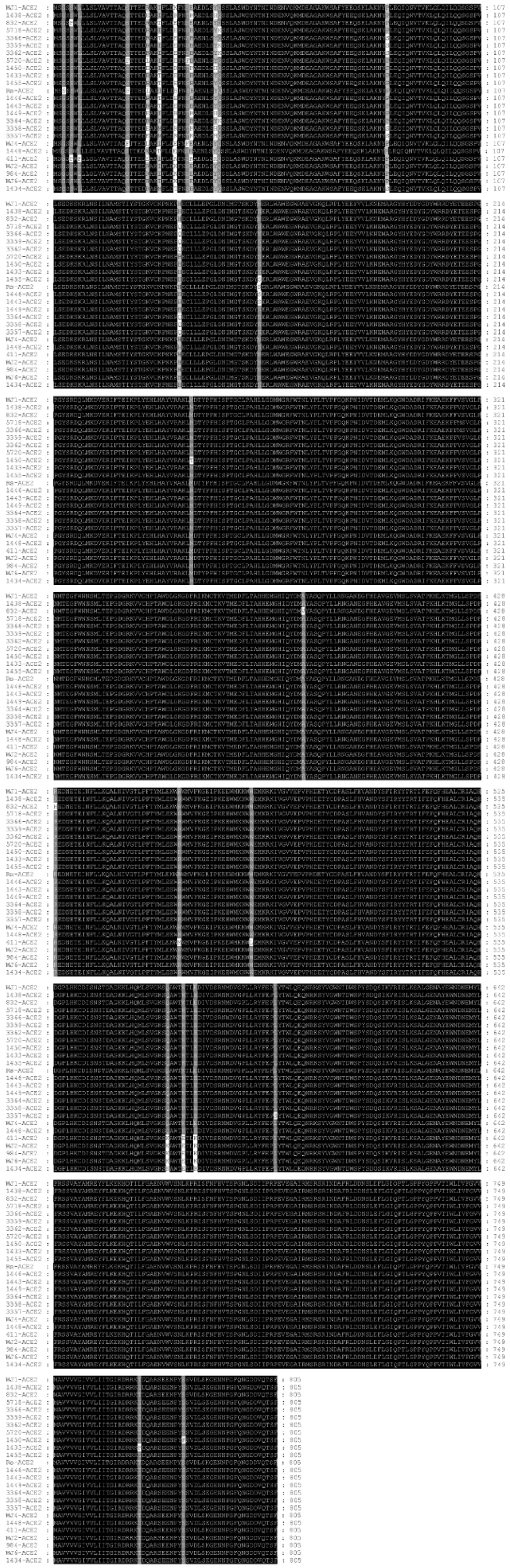

25

**Fig. S2. Analysis of different *R.sinicus* ACE2s usage of SARSr-CoVs determined by immunofluorescence assay.** (A-D) Determination of bat SARSr-CoVs infectivity in HeLa cells with and without the expression of ACE2 from *R. sinicus* or human (hACE2) at an MOI=1. ACE2 expression was detected with mouse anti-Stag monoclonal antibody followed by FITC-labelled goat anti-mouse IgG H＆L. At 24h after infection, virus replication was detected using rabbit serum against the SARSr-CoV-Rp3 Np followed by Cy3-conjugated goat anti-rabbit IgG. Nuclei were stained with DAPI. (A) SARSr-CoV-RsWIV1. (B) SARSr-CoV-RsWIV16. (C) SARSr-CoV-Rs4231. (D) SARSr-CoV-RsSHC014.

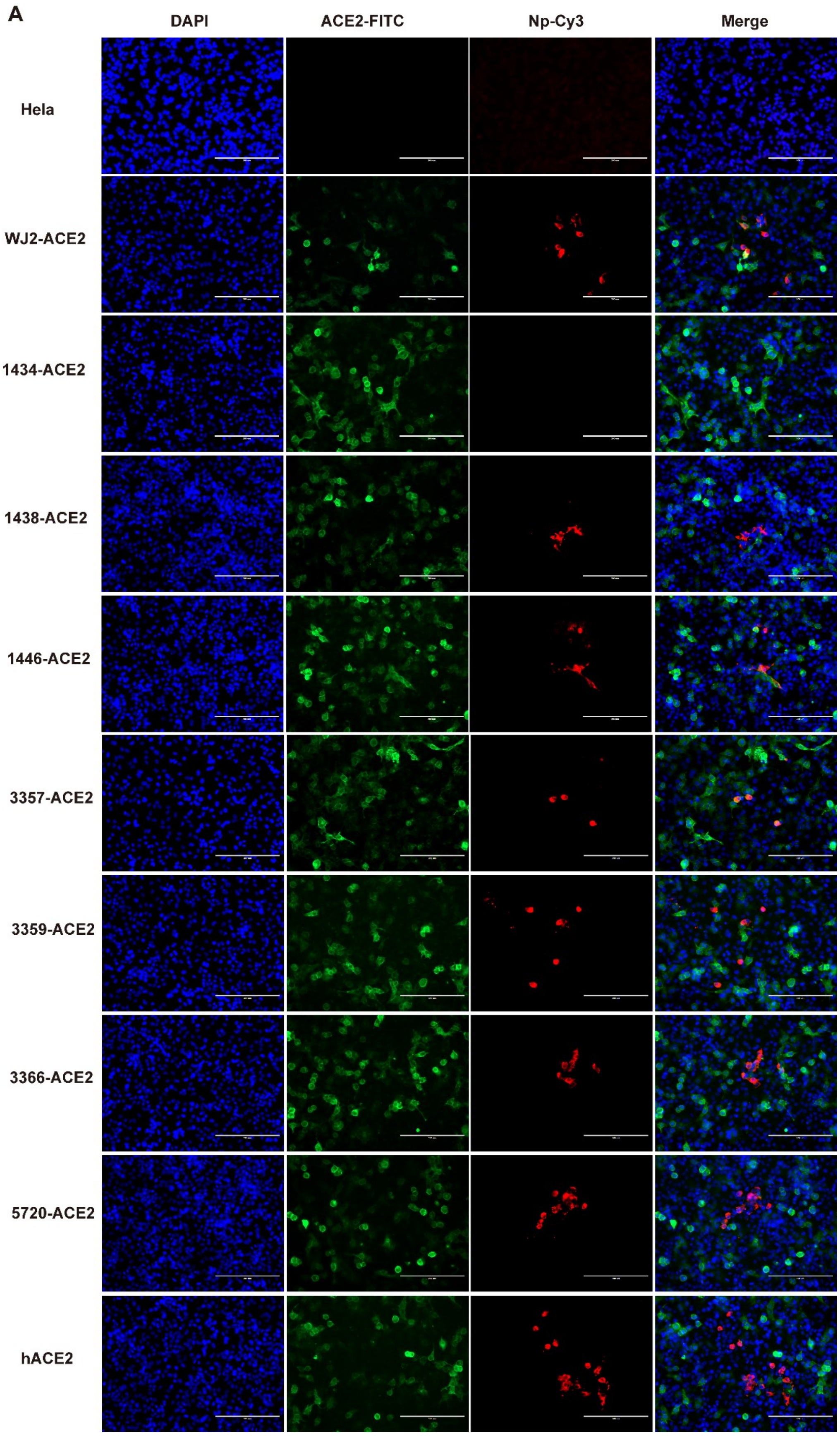
A
DAPI
ACE2-FITC
Np-Cy3
Merge
Hela
WJ2-ACE2
1434-ACE2
1438-ACE2
1446-ACE2
3357-ACE2
3359-ACE2
3366-ACE2
5720-ACE2
hACE2

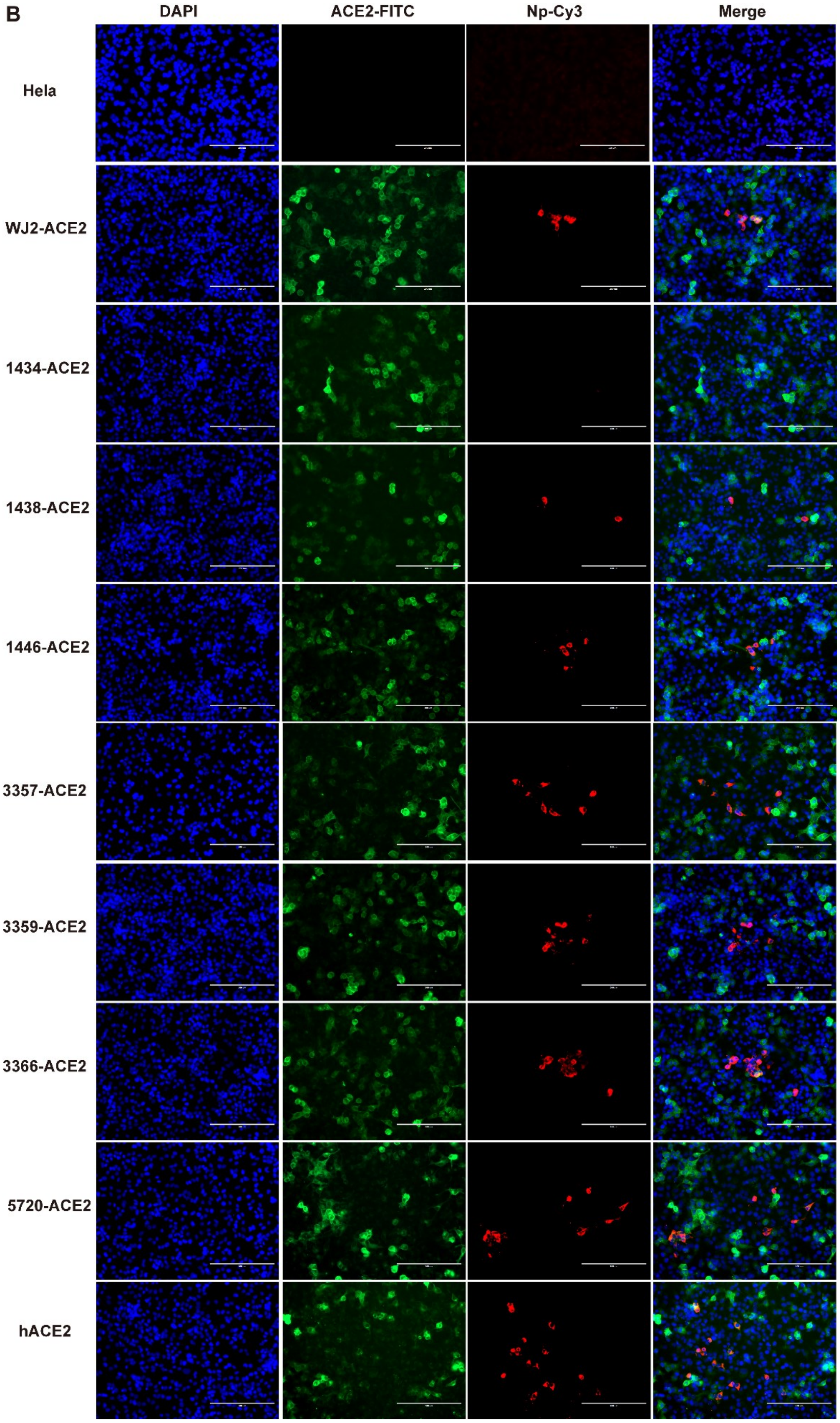
B
DAPI
ACE2-FITC
Np-Cy3
Merge
Hela
WJ2-ACE2
1434-ACE2
1438-ACE2
1446-ACE2
3357-ACE2
3359-ACE2
3366-ACE2
5720-ACE2
hACE2

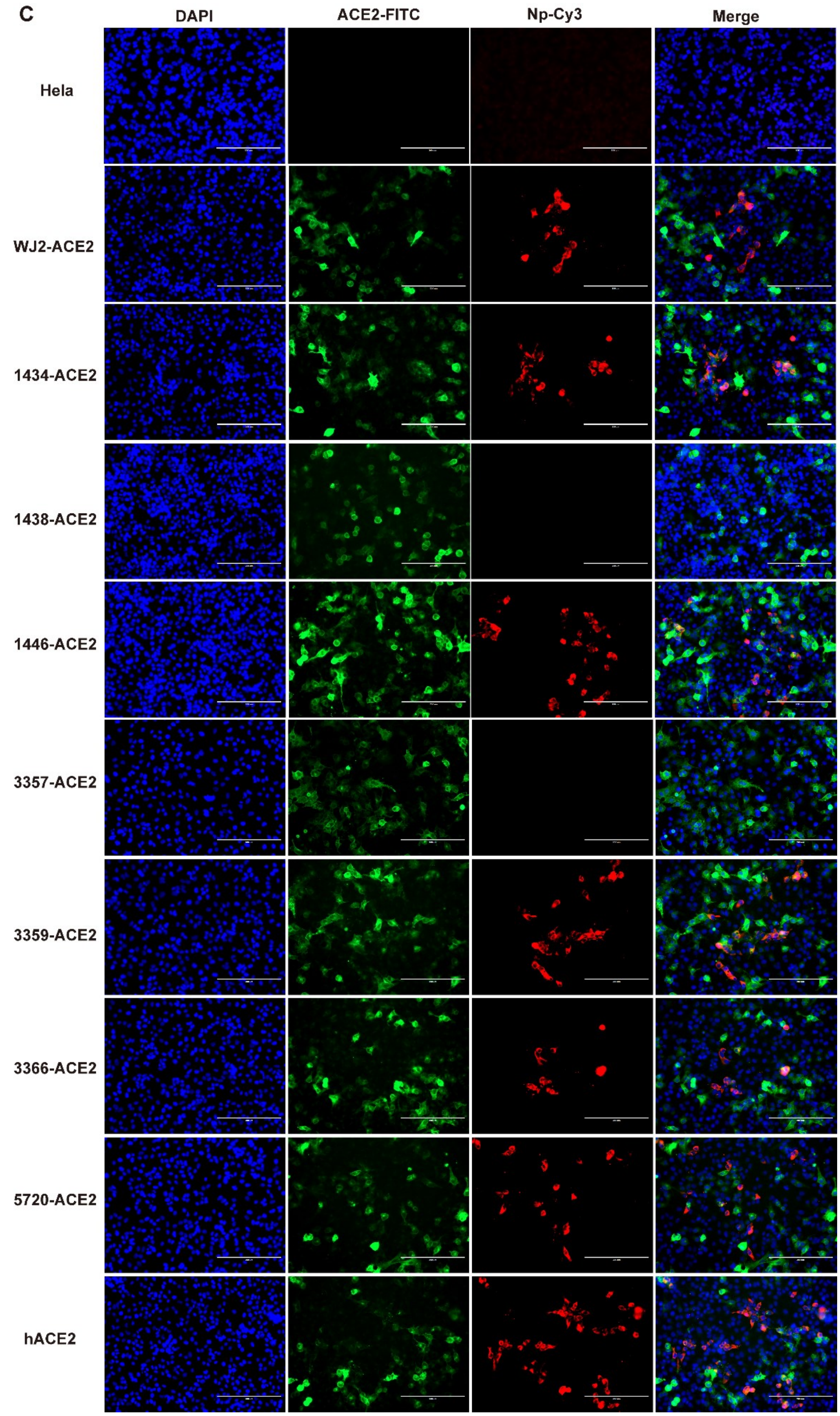
C
DAPI
ACE2-FITC
Np-Cy3
Merge
Hela
WJ2-ACE2
1434-ACE2
1438-ACE2
1446-ACE2
3357-ACE2
3359-ACE2
3366-ACE2
5720-ACE2
hACE2

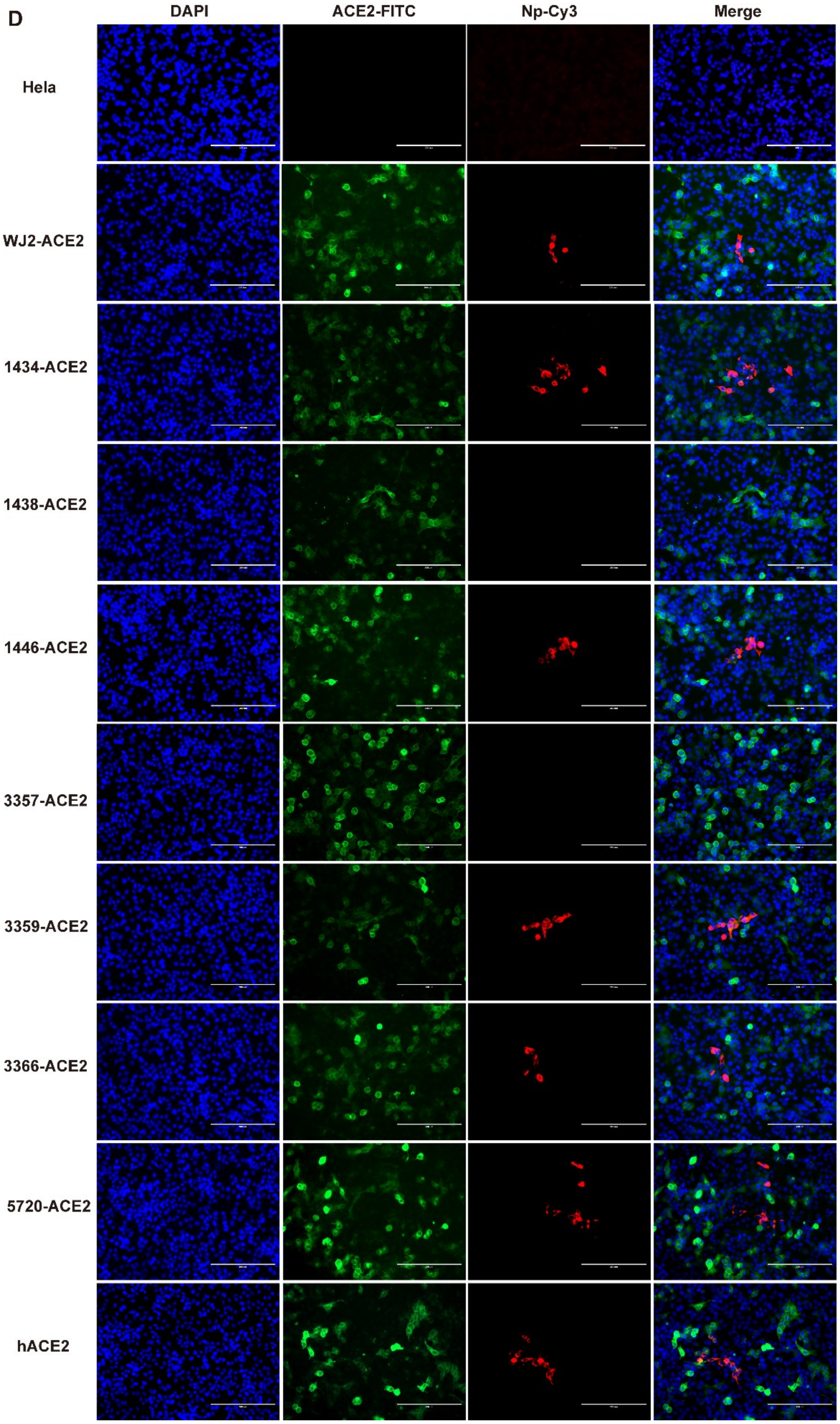
D
DAPI
ACE2-FITC
Np-Cy3
Merge
Hela
WJ2-ACE2
1434-ACE2
1438-ACE2
1446-ACE2
3357-ACE2
3359-ACE2
3366-ACE2
5720-ACE2
hACE2

**Fig. S3. The expression of ACE2 in HeLa cells in SARS-CoV BJ01 pseudotyped infection assay.** ACE2 expression was detected with mouse anti-Stag monoclonal antibody followed by HRP-labelled goat anti-mouse IgG antibody. β-actin was detected with mouse anti-β-action monoclonal antibody by HRP-labelled goat anti-mouse IgG antibody.

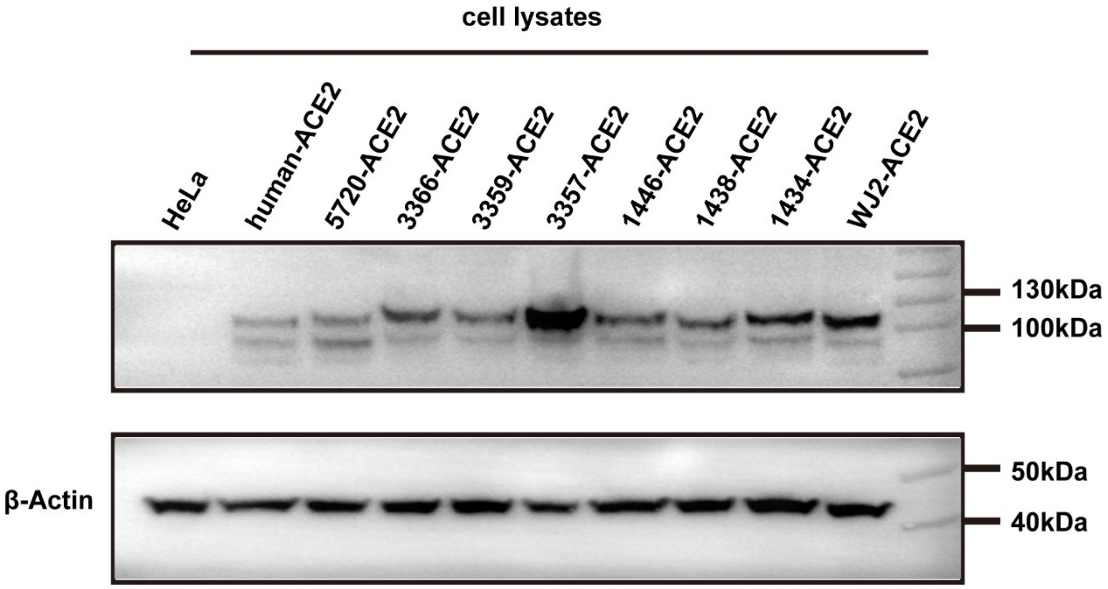

**Fig. S4. The saturation profiles for the indicated binding pair measured by Octet RED system.** (A-C) Binding assay of human ACE2 or bat ACE2 to WIV1 RBD. (D-F) Binding assay of human ACE2 or bat ACE2 to SHC014 RBD. (G-I) Binding assay of human ACE2 or bat ACE2 to BJ01 RBD. Different RBD proteins were immobilized on the sensors and tested for binding with gradient concentrations of *R.sinicus* ACE2s, hACE2 or hDPP4. And the Y axis shown the real-time binding response. The coefficient of determination ($R^2$) for these interactions was close to 1.0. The profiles of RsWIV1-RBD/*R.sinicus* ACE2-1434, RsSHC014-RBD/*R.sinicus* ACE2-3357, and BJ01-RBD/*R.sinicus* ACE2-3357 binding affinity didn't show because there were not obviously binding activity between them.

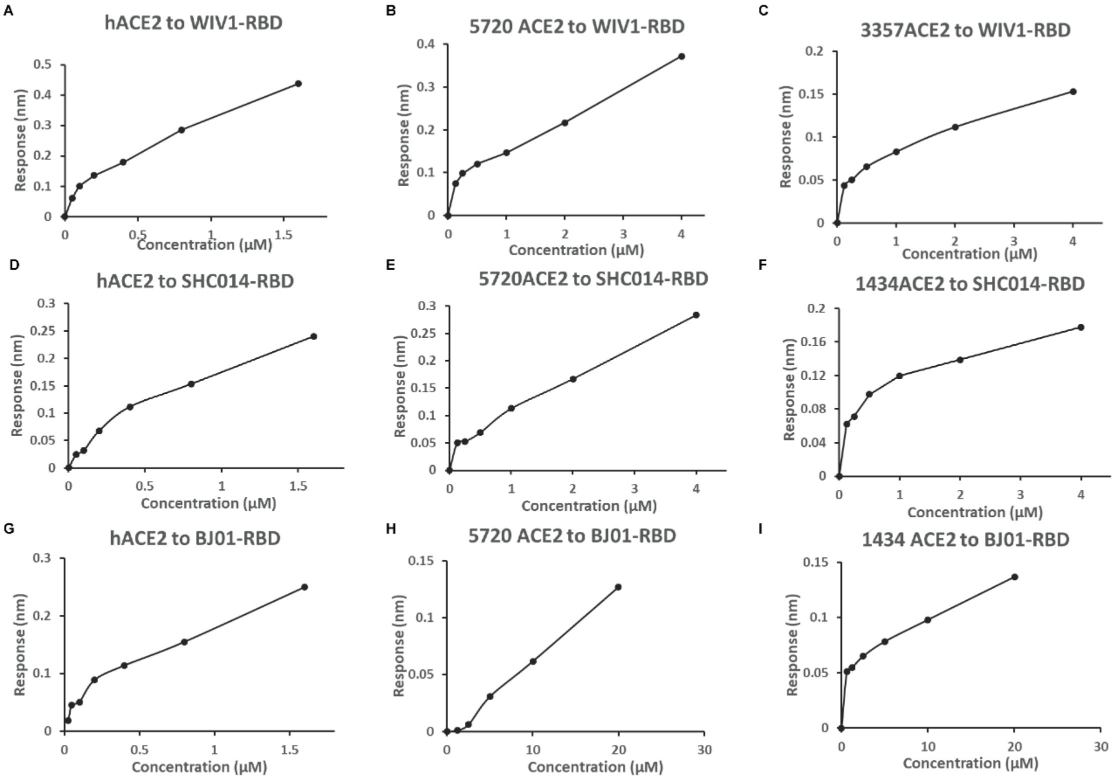

**Fig. S5. The positive selection residues in *R.sinicus* ACE2 and SARSr-CoV RBD.** Eight residues under positive selection (red) in *R.sinicus* ACE2 and five residues under positive selection in SARSr-CoV RBD (blue) are shown in the co-crystal structure of human ACE2 and SARS-CoV RBD (PDB 2AFJ). Human ACE2 is in cyan and SARS-CoV RBD is in deep pink.

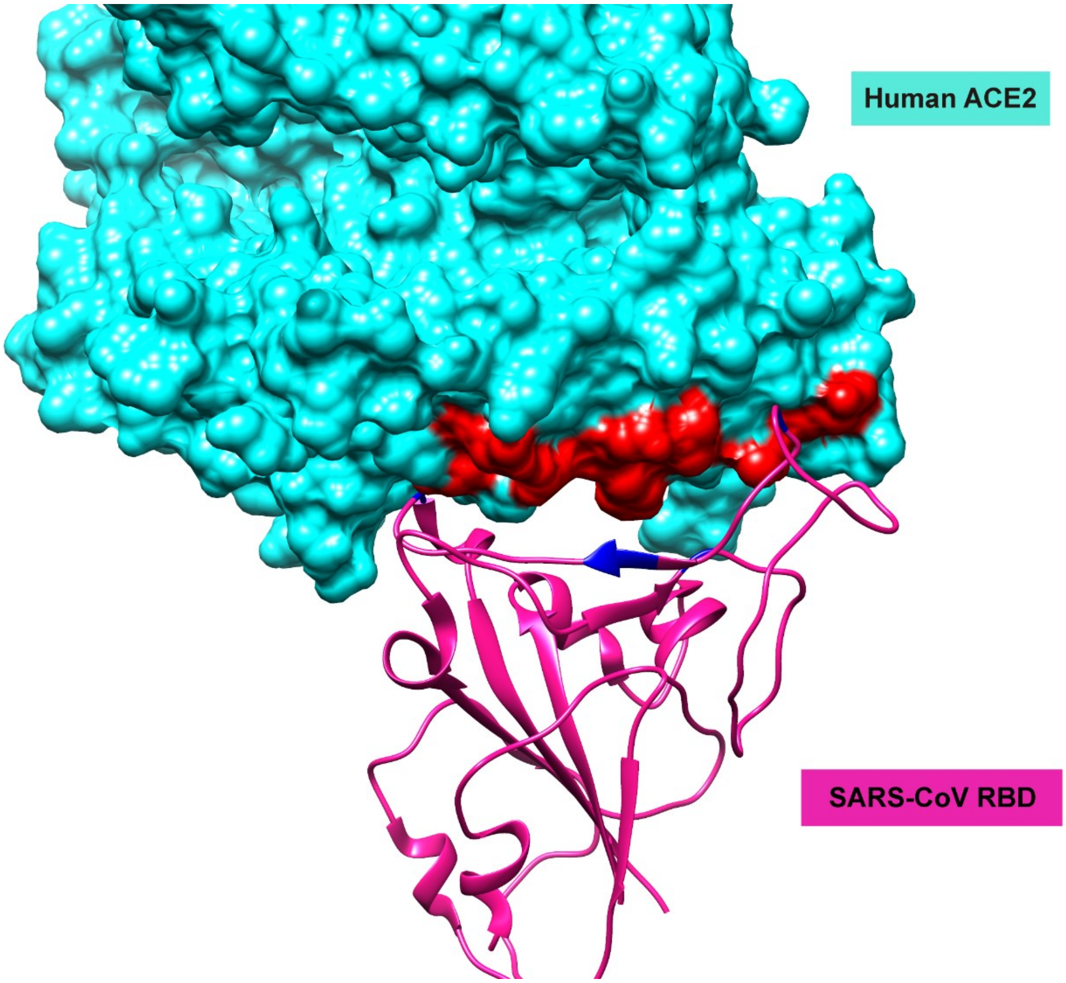

**Fig. S6. Alignment of amino acids sequences of *R. affinis* ACE2.** Residues 1-804 are shown. The accession numbers of various sequences used in this alignment are list in Table S2.

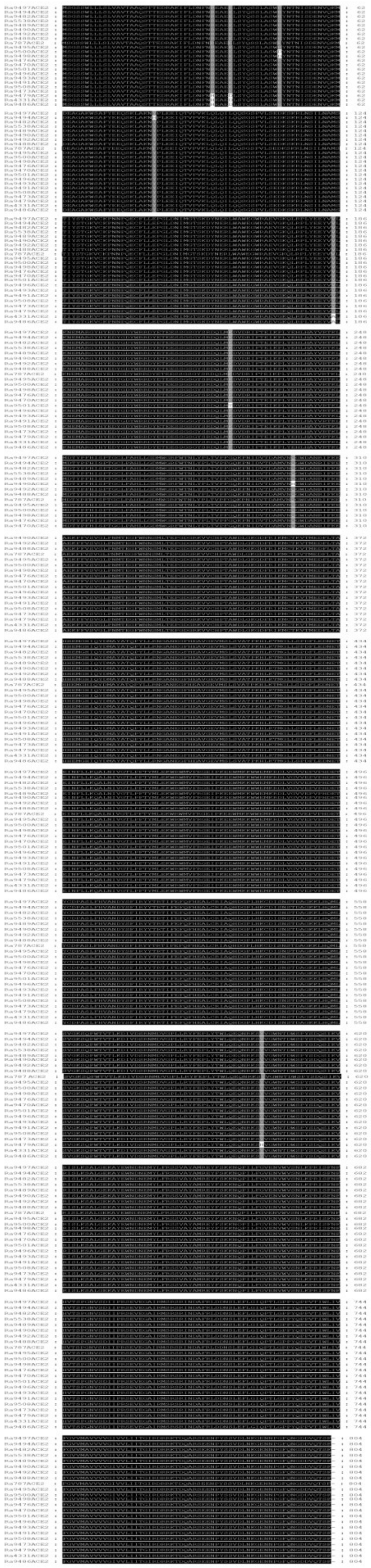

**Table S2. Database accession number of sequences used in this study.**

| Dataset | Sources | Sample ID | Accession number |
|---|---|---|---|
| *Rhinolophus sinicus* ACE2 | GenBank database | *R.s*-832 | GQ999936 |
| | GenBank database | *R.s*-411 | GQ999933 |
| | GenBank database | *R.s*-ACE2 | GQ262791 |
| | GenBank database | *R.s*-3357 | KC881004 |
| | Obtained in this study | *R.s*-WJ4 to 3362 | MT394181-201 |
| *Rhinolophus affinis* ACE2 | Obtained in this study | *R.a*-787 to 4331 | MT394203-225 |
| SARSr-CoV | GenBank database | SARSr-CoV-RsWIV16 | KT444582 |
| | | SARSr-CoV-RsWIV1 | KF367457 |
| | | SARSr-CoV-Rs3367 | KC881006 |
| | | SARSr-CoV-Rs4084 | KY417144 |
| | | SARSr-CoV-RsSHC014 | KC881005 |
| | | SARSr-CoV-Rs7327 | KY417151 |
| | | SARSr-CoV-Rs4231 | KY417146 |
| | | SARSr-CoV-Rs4874 | KY417150 |
| | | SARSr-CoV-Rs9401 | KY417152 |

**Table S3**. **The mutations in the RBMs of various SARSr-CoV strains.** Five residues directly contact with ACE2 are shown. The accession numbers of various sequences used in this table see Table S2

| Strains | Host | 442 | 472 | 479 | 480 | 487 |
|---|---|---|---|---|---|---|
| hBJ01 | Human | Y | L | N | D | T |
| RsWIV1 | *R.sinicus* | S | F | N | D | N |
| RsWIV16 | *R.sinicus* | S | F | N | D | N |
| Rs4231 | *R.sinicus* | W | P | R | P | A |
| Rs3367 | *R.sinicus* | S | F | N | D | N |
| Rs7327 | *R.sinicus* | S | F | N | D | N |
| Rs4084 | *R.sinicus* | W | P | R | P | A |
| Rs4874 | *R.sinicus* | S | F | N | D | N |
| Rs9401 | *R.sinicus* | S | F | N | D | N |
